## Supplementary material for "Charged peptides enriched in aromatic residues decelerate condensate ageing driven by cross-β-sheet formation": Suppementary Material

### SI. CALVADOS2 MODEL DETAILS

The coarse-grained CALVADOS2 force field [1] is given by:

$$u_{\mathrm{CALVADOS2}} = u_{\mathrm{Bonds}} + u_{\mathrm{Electrostatic}} + u_{\mathrm{Hydrophobic}}, \tag{S1}$$

where $u_{\mathrm{Hydrophobic}}$ and $u_{\mathrm{Electrostatic}}$ interactions are only applied between non-bonded beads and $u_{\mathrm{Bonds}}$ between subsequent beads directly bonded to each other.

Bonded interactions between subsequent amino acid protein beads are described by an harmonic potential:

$$u_{\mathrm{Bonds}} = k_b(r - r_0)^2, \tag{S2}$$

where the equilibrium bond length is $r_0 = 3.8$ Å between bonded amino acid beads. The spring constant is $k_b = 9.6$ kcal mol$^{-1}$ Å$^{-2}$.

The hydrophobic interactions between different amino acid types are implemented through the functional form of an Ashbaugh/Hatch potential [2, 3]:

$$u_{\mathrm{Hydrophobic}} = \begin{cases} 4\epsilon_{ij}\left[\left(\frac{\sigma_{ij}}{r}\right)^{12} - \left(\frac{\sigma_{ij}}{r}\right)^{6}\right] + (1-\lambda_{ij})\epsilon_{ij}, & r < 2^{1/6}\sigma_{ij} \\ \lambda_{ij}4\epsilon_{ij}\left[\left(\frac{\sigma_{ij}}{r}\right)^{12} - \left(\frac{\sigma_{ij}}{r}\right)^{6}\right], & \text{otherwise,} \end{cases} \tag{S3}$$

where $\lambda_i$ and $\lambda_j$ are parameters that account for the hydrophobicity of the ith and jth interacting particles respectively, being $\lambda_{ij} = (\lambda_i + \lambda_j)/2$. The excluded volume (i.e. molecular diameter) of the different residues is given by $\sigma_i$ and $\sigma_j$, where $\sigma_{ij} = (\sigma_i + \sigma_j)/2$, and r is the distance between the ij particles. $\epsilon_{ij}$ is set to 0.2 kcal/mol [4]. The specific values for each amino acid $\sigma$, q, and $\lambda$ parameters can be found in reference [1].

The electrostatic interactions, $u_{\mathrm{Electrostatic}}$, among charged amino acids is described by a Coulomb/Debye-Hückel potential of the form:

$$u_{\mathrm{Electrostatic}} = \frac{1}{4\pi D}\frac{q_i q_j}{r}e^{-r/\kappa}, \tag{S4}$$

where $q_i$ and $q_j$ represent the charges of the beads i and j. The $u_{\mathrm{Electrostatic}}$ term permits to avoid Ewald summation and screening of electrostatic interactions by the implict solvent and ions. Within this model the Debye length ($\kappa$) is not a fixed value, but rather depends on the salt concentration, allowing for a better description of LLPS modulation with varying salt concentration. The Debye length is defined as:

$$\kappa = \sqrt{\frac{1}{8\pi B c_s}} \tag{S5}$$

where $c_s$ is the ionic strength and B is the Bjerrum length, defined as:

$$B = \frac{1.671x10^{-5}}{DT}. \tag{S6}$$

In this model, the relative dielectric constant of the solvent (D) follows the empirical equation [5]:

$$D = \frac{5321}{T} + 233.76 - 0.9297T + 1.417x10^{-3}T^2 - 8.292x10^{-7}T^3. \tag{S7}$$

$u_{Electrostatic}$ and $u_{Hydrophobic}$ are truncated at 40 and 20 Å respectively, as recommended in reference [1]. The timestep for the integration of the equations of motion is 10 fs. The relaxation time for the Nose-Hoover [6, 7] thermostat and barostat thermostat and barostat is 5 ps. Periodic boundary conditions are applied in the 3 directions of space.

### SII. NPT SIMULATIONS

As outlined in the main text we compute the critical temperature for phase separation for the different protein-peptide systems through NPT simulations. In this manner, we compute the condensed phase branch of the temperature-density phase diagram (Figure 2(a) in main text). The critical temperature is determined as the average between the last temperature at which the condensate remains liquid and the lowest one at which the condensate disassembles. It must be noted that we compute the phase diagram in this approximate way, opposed to using the Direct Coexistence method [8] in order to have a better control of the condensate concentration, opposed to tuning the total system composition. This NPT method is approximate, since we assume that the vapor pressure is equal to 0 at all temperatures, which can lead to an underestimation of the critical temperature. We verified that this approach is valid with protein/RNA mixtures in Ref. [9].

### SIII. AGING ALGORITHM

We perform our aging simulations allowing the formation of inter-protein β-sheets to take place, according to the scheme developed in Refs. [10–12]. We first identified the regions of the proteins sequence that are prone to forming inter-protein secondary structures, as reported in references [13, 14]. These are low-complexity aromatic-rich kinked segments (LARKS). In the FUS LCD sequence we consider the following: $_{37}$SYSGYS$_{42}$ (PDB code 6BWZ), $_{54}$SYSSYGQS$_{61}$ (PDB code 6BXV), and $_{77}$STGGYG$_{82}$ (PDB code 6BZP) [13], while for TDP-43 LCD we consider $_{27}$GNNQGS$_{32}$ (PDB 5WKD), $_{39}$NFGAFS$_{44}$ (PDB 5WHN), $_{55}$AALQSS$_{60}$ (PDB 6CB9), $_{60}$SWGMMGMLASQ$_{70}$ (PDB 6CFH), and $_{123}$GFNGGFG$_{139}$ (PDB 5WIQ) [14]. In Ref. [10] the potential of mean force of the structured and disordered conformations of such regions of FUS LCD was calculated, indicating us what the interaction difference between before and after such structure is formed is. Similar calculations were performed in Ref. [15] for TDP-43 LARKS. Our aging simulations consist of non-equilibrium Molecular Dynamics runs in which, when the aforementioned regions are found in space, an effective change in the pairwise interaction of the involved residues (according to the PMF calculations) is performed. We use a distance criterion, in which at least the central residues of four LARKS of the same kind must meet within a cutoff distance of 15.5 Å for FUS LCD and 13 Å for TDP-43 LCD. This distance is chosen so that the formation of the inter-protein β-sheets takes place within an accessible timescale. Moreover, the requirement of at least four LARKS of different protein replicas meeting simultaneously is set so that the structures reported in Ref. [13] can be sustained. In this way we model the formation of inter-protein β-sheets. This is done with the *fix bond/react* command [16] available in LAMMPS.

Once the disorder-to-order structural transition takes place, we also impose an harmonic potential between directly bonded residues to account for the rigidity of the amino acids involved in the cross-β-sheet motif. This potential follows the equation

$$u_{Angular} = k_a(\theta - \theta_0), \tag{S8}$$

and applies to the angle formed between all the directly-bonded residues in the cross-β-sheet formed. In our simulations, $k_a$ is 5 kcal mol$^{-1}$ rad$^{-2}$, $\theta$ is the angle formed by three consecutive residues and $\theta_0$ is the equilibrium value of the angle, set to 180°. We perform the ageing simulations in the canonical ensemble, fixing the equilibrium density of the condensate, which is obtained from the NPT simulations. Example files of this algorithm can be found in the online repository https://doi.org/10.5281/zenodo.15168499 , where we have included configuration files for both FUS LCD and TDP-43 LCD, as well as the LAMMPS scripts and associated files needed to run a non-equilibrium simulation.

With our computational setup, we are capable of performing simulations at a speed of 600 timesteps per CPU second, which allows us to simulate approximately 500 ns per day for systems of $\sim$15000 particles (i.e., $\sim$ 100 protein replicas) and 64 CPUs per simulation (tested with Intel Xeon Platinum 8160 CPU).

### SIV. G(T), G' AND G" CALCULATION

In our simulations, the storage modulus (G') and loss modulus (G") are obtained from the stress relaxation modulus G(t), which describes how stress relaxes over time after an initial perturbation. G(t) is obtained from the stress autocorrelation function via the Green-Kubo relation:

$$\eta = \int_0^\infty dt\, G(t) \tag{S9}$$

In an isotropic system, we can compute the shear relaxation modulus $G(t)$ more accurately by using all the components of the pressure tensor ($\sigma_{\alpha\beta}$) as shown in Ref. [17]:

$$\begin{aligned} G(t) = {} & \frac{V}{5k_BT}\left[\langle\sigma_{xy}(0)\sigma_{xy}(t)\rangle + \langle\sigma_{xz}(0)\sigma_{xz}(t)\rangle + \langle\sigma_{yz}(0)\sigma_{yz}(t)\rangle\right] \\ & + \frac{V}{30k_BT}\left[\langle N_{xy}(0)N_{xy}(t)\rangle + \langle N_{xz}(0)N_{xz}(t)\rangle + \langle N_{yz}(0)N_{yz}(t)\rangle\right], \end{aligned} \tag{S10}$$

where $N_{\alpha\beta} = \sigma_{\alpha\alpha} - \sigma_{\beta\beta}$ is the first normal stress difference. This correlation can be easily computed by using the compute ave/correlate/long in the USER-MISC package of LAMMPS [18] or EXTRA-FIX in LAMMPS versions after 2021. The storage $G'(\omega)$ and loss $G''(\omega)$ moduli can be directly calculated from the complex modulus $G^*(\omega)$ defined as

$$G^*(\omega) = G'(\omega) + iG''(\omega). \tag{S11}$$

The complex modulus is calculated as the Laplace transform of $G(t)$ and thus, the storage and loss moduli can be computed as the sine and cosine transforms of $G(t)$, i.e.:

$$G'(\omega) = \omega \int_0^\infty dt\, G(t)\sin(\omega t), \tag{S12}$$

$$G''(\omega) = \omega \int_0^\infty dt\, G(t)\cos(\omega t), \tag{S13}$$

Please see [19] for further details. These equations relate the time-dependent stress relaxation function to the material's frequency-dependent viscoelastic response. In this context, the frequency ($\omega$) refers to the angular frequency of an oscillatory deformation that the system would experience in a rheological experiment. In experimental rheology, an oscillatory shear is applied, and the material's response is measured. This approach allows MD simulations to predict rheological behavior without explicitly running large-scale oscillatory deformation simulations.

In Figure S1 we show the time evolution of G' and G" with time as inter-protein β-sheets accumulate in the bulk FUS LCD condensate. The crossover between these two quantities at low frequencies ($<10^9$ rad/s) indicate the transition into a solid-like behaviour of the condensate, and the formation of a percolated network of cross-β-sheets.

Moreover, the viscosity can be directly computed from $G(t)$. At short times, $G(t)$ is smooth and the integral can be computed using numerical integration (trapezoidal rule). However, at longer times $G(t)$ presents more noise, and hence, we calculate the integral in that regime by first fitting $G(t)$ to a series of Maxwell modes ($G_i \exp(-t/\tau)$) equidistant in logarithmic time [20] and then by calculating the integral analytically. Our fit to the Maxwell modes is carried out with the help of the open-source RepTate software (version 1.1.1 20200602) [21]. Finally, viscosity is obtained by adding the two terms:

$$\eta = \eta(t_0) + \int_{t_0}^\infty dt\, G_M(t), \tag{S14}$$

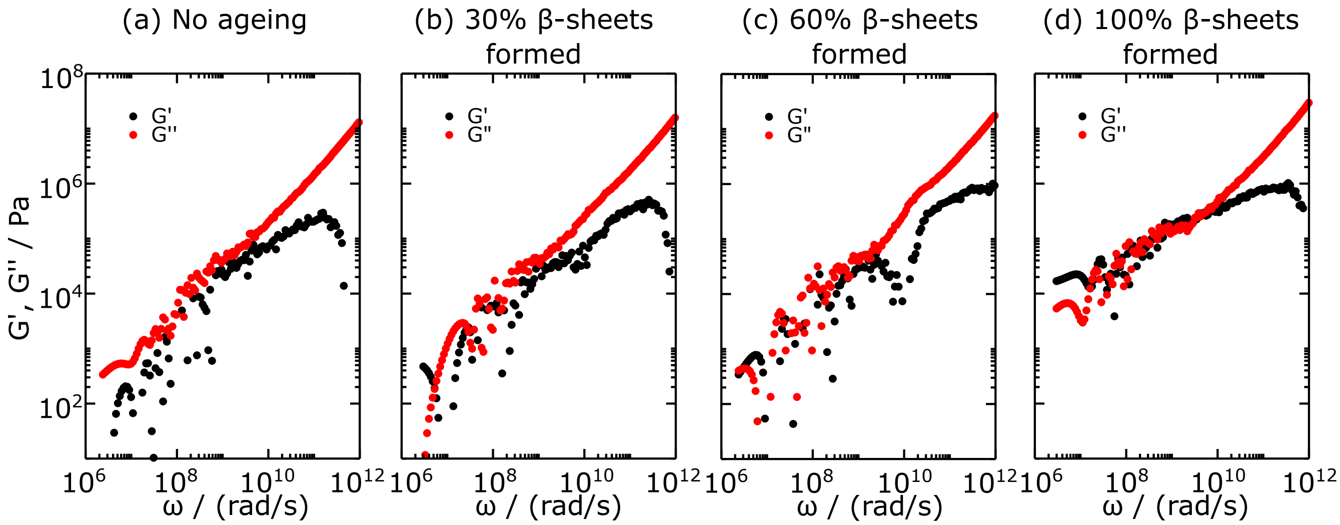


Figure S1: (a)-(d) Storage (G') and shear loss (G") moduli as a function of frequency (ω) for FUS LCD condensates at different stages of the ageing process.

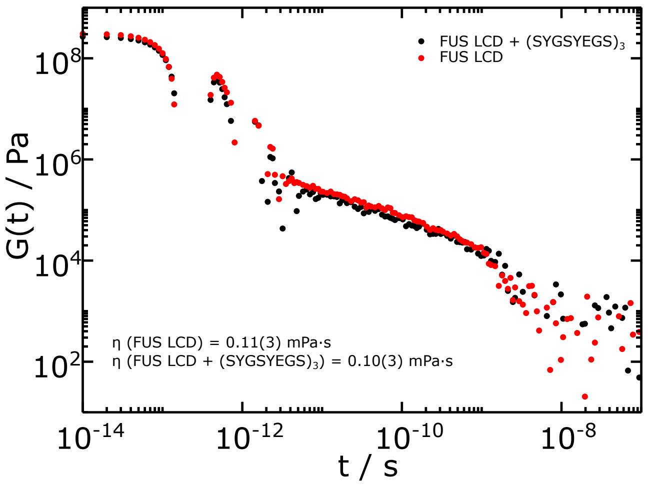


Figure S2: Shear stress relaxation modulus (G(t)) of the FUS LCD bulk condensed phase in presence and absence of the $(SYGSYEGS)_3$.

where $\eta(t_0)$ corresponds to the computed term for short time-scales, $G_M(t)$ is the part evaluated via the Maxwell modes fit at long time-scales, and $t_0$ is the time that separates both.

In Figure S2 we show $G(t)$ for FUS LCD and FUS LCD in presence of the $(SYGSYEGS)_3$ at a concentration of 0.07 mg $(SYGSYEGS)_3$/mg FUS LCD. The resemblance between the two curves, as well as the calculation of the viscosity as described above (0.11(3) and 0.10(3) mPa·s in absence and presence of the small peptide respectively) proves that the addition of the small peptides in very moderate concentrations has a rather small impact in the viscoelastic properties of the protein condensates.

### SV. OBTAINING PHASE DIAGRAMS VIA DIRECT COEXISTENCE SIMULATIONS

When the phase diagram is calculated via the Direct Coexistence method, the critical point of the phase diagrams is estimated using the universal scaling law of coexistence densities near a critical point [22], and the law of rectilinear diameters [23]:

$$\left(\rho_l(T) - \rho_v(T)\right)^{3.06} = d\left(1 - \frac{T}{T_c}\right) \quad \text{(S15)}$$

and

$$(\rho_l(T) + \rho_v(T))/2 = \rho_c + s_2(T_c - T) \quad \text{(S16)}$$

where $\rho_l$ and $\rho_v$ refer to the coexisting densities of the condensed and diluted phases respectively, $\rho_c$ is the critical density, $T_c$ is the critical temperature, and $d$ and $s_2$ are fitting parameters.

### SVI. AGGREGATION CURVES

In this section we provide all the aggregation curves for all of the systems studied, as detailed in the main text. All of the FUS LCD systems are reported in Figure S3, while those for TDP-43 LCD are reported in Figure S4. In all cases, our simulations were carried out with 100 protein replicas and varying amounts of the peptides, as indicated for each case. Moreover, in Table S1 we report all of the peptide concentrations at which the ageing simulations reported in the main text and Figures S3 and S4 were performed. For these concentrations we found that the critical temperature decreases about $\sim$ 0.03 $T/T_c$, as indicated in Figure 2(b) of the main text.

We shall note that for the FUS + $(SYGESYGSYGEG)_2$ and TDP-43 LCD + $(SYGSYEGS)_3$ there is one trajectory in which we have not observed aggregation in each system after over 3000 and 4000 ns respectively. Due to computational limitations, we have estimated the nucleation time (represented in Figure S5), the aggregation half time (represented in Figure 3 of the main text) and the primary nucleation rate ($k_n$) (represented in Figure S6) based on the assumption that the aggregation process is dominated by the elongation step, and limited by the nucleation step, as explained in the main text. In that way, we approximate the nucleation events from the Poisson distribution formula for a number of events that occur after a given observation time: $N(t) = N_0 exp(-\lambda t)$, where N is the non-nucleated seeds, $N_0$ is the initial number of seeds, λ is a constant and t is the time. With the data available for the seeds that have nucleated, we can obtain λ and later calculate the estimated time that it will take for the remaining seed to nucleate.

### SVII. SIMULATION CONCENTRATION AND DENSITY DETAILS

In table S1 we summarize the concentration and density used for the ageing simulations.

| Guest molecule | FUS LCD | | TDP-43 LCD | |
|---|---|---|---|---|
| | mg Guest Molecule/ mg Protein | Condensate density / (g cm$^{-3}$) | mg Guest Molecule/ mg Protein | Condensate density / (g cm$^{-3}$) |
| Protein only | - | 0.334 | - | 0.352 |
| + $(SYSYKRKS)_3$ | 0.042 | 0.310 | - | - |
| + $(SYSYKRKK)_3$ | 0.044 | 0.309 | 0.055 | 0.306 |
| + $(SYGSYEGS)_3$ | 0.070 | 0.320 | 0.131 | 0.320 |
| + $(SYGESYGSYGEG)_2$ | 0.069 | 0.322 | 0.130 | 0.320 |
| + $(SYGSYGSYEGEG)_2$ | 0.069 | 0.322 | 0.130 | 0.320 |
| + $(SYGKSYGSYGKG)_2$ | - | - | 0.087 | 0.314 |

Table S1: Small peptide concentration used in the ageing simulations

### SVIII. KINETIC CONSTANT AND NUCLEATION TIME ANALYSIS

We perform multiple kinetic analysis of our aggregation curves. In Figure 3 (main text) we show the half time ($t_{1/2}$), defined as the point where the relative aggregate concentration value is halfway between the initial value and final plateau value. In Figure S5 we show the nucleation time, calculated as the time it takes for the first β-sheet to be formed. Moreover, in Figure S6 we show the primary nucleation rate ($k_n$), obtained from Amlylofit fitter [24]. For this last analysis

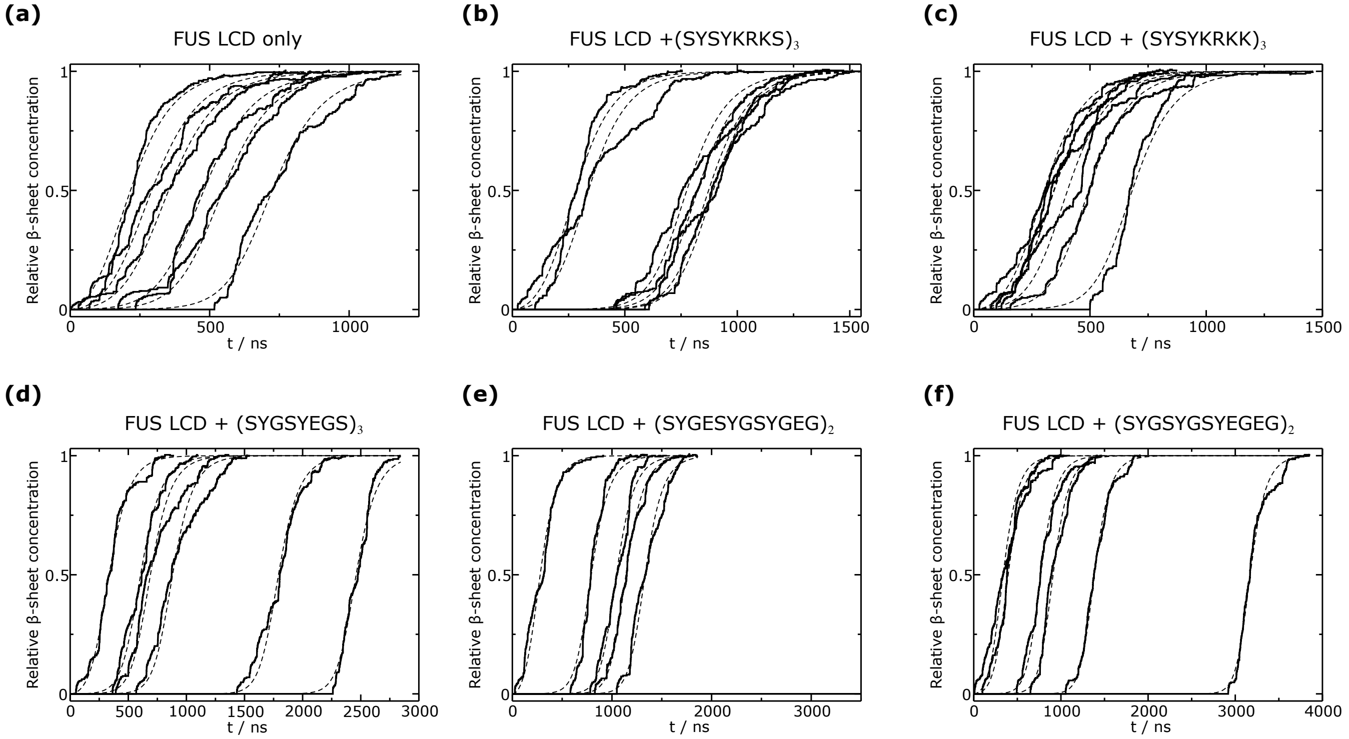


Figure S3: (a)-(f) Relative aggregate concentration for all the FUS LCD systems, as indicated for each of them. For each system, the different curves represent independent trajectories. The dashed lines indicate kinetic fits performed with Amylofit software [24]. For the nature of this analysis see Section SVIII.

| Guest molecule | Charge | # Aromatic residues |
|---|---|---|
| $(SYSYKRKS)_3$ | +6 | 6 |
| $(SYSYKRKK)_3$ | +9 | 6 |
| $(SYGSYEGS)_3$ | -3 | 6 |
| $(SYGESYGSYGEG)_2$ | -4 | 6 |
| $(SYGSYGSYEGEG)_2$ | -4 | 6 |
| $(SYGKSYGSYGKG)_2$ | +4 | 6 |

Table S2: Charge and aromatic residue count for all the small peptides considered in Figure 3 & 4 (main text).

we followed a particular approach: We first fit all of the nucleation curves independently using the secondary nucleation dominated mechanism, as outlined in the main text. From this analysis, we obtain 3 kinetic constants: (1) the primary nucleation rate $k_n$, the rate at which primary nuclei are formed, (2) $k_2$, the secondary nucleation rate constant, and (3) $k_+$, the elongation rate constant. Since the primary nucleation is the limiting and differentiating stage in the aggregation in our simulations, we then average the values of $k_2$ and $k_+$ for all systems of the same protein (either FUS LCD or TDP-43 LCD) and repeat the kinetic fits using such averaged values. This procedure allows us to directly compare the kinetic constant of the differentiating step of the aggregation process, to determine whether ageing takes place faster or slower for the different systems. We plot the obtained values of -log($k_n$) as box plots in Figure S6. By comparing the box plots from Figure S6 with those from Figure S5 and Figure 3 (main text) we can conclude that all the different analysis yield equivalent conclusions regarding the deceleration of ageing for certain systems, as discussed in the main text.

Lastly, we perform a Mann–Whitney U test to determine whether the condensates with peptides have a statistically different nucleation half-time (reported in Figure 3 of the main text). In table S3 we summarize the U and p values that determine whether the formation of β-sheets takes place at a statistically different pace. For 6 seeds, we perform a 1-tailed analysis with a p values of 0.05, for which the U threshold value is 7, meaning that U values equal or lower than 7 and p values lower than 0.05 represent that the two systems are statistically different in the β-sheet formation process. We find that only FUS LCD+$(SYSYKRKK)_3$ is statistically alike to FUS LCD only.

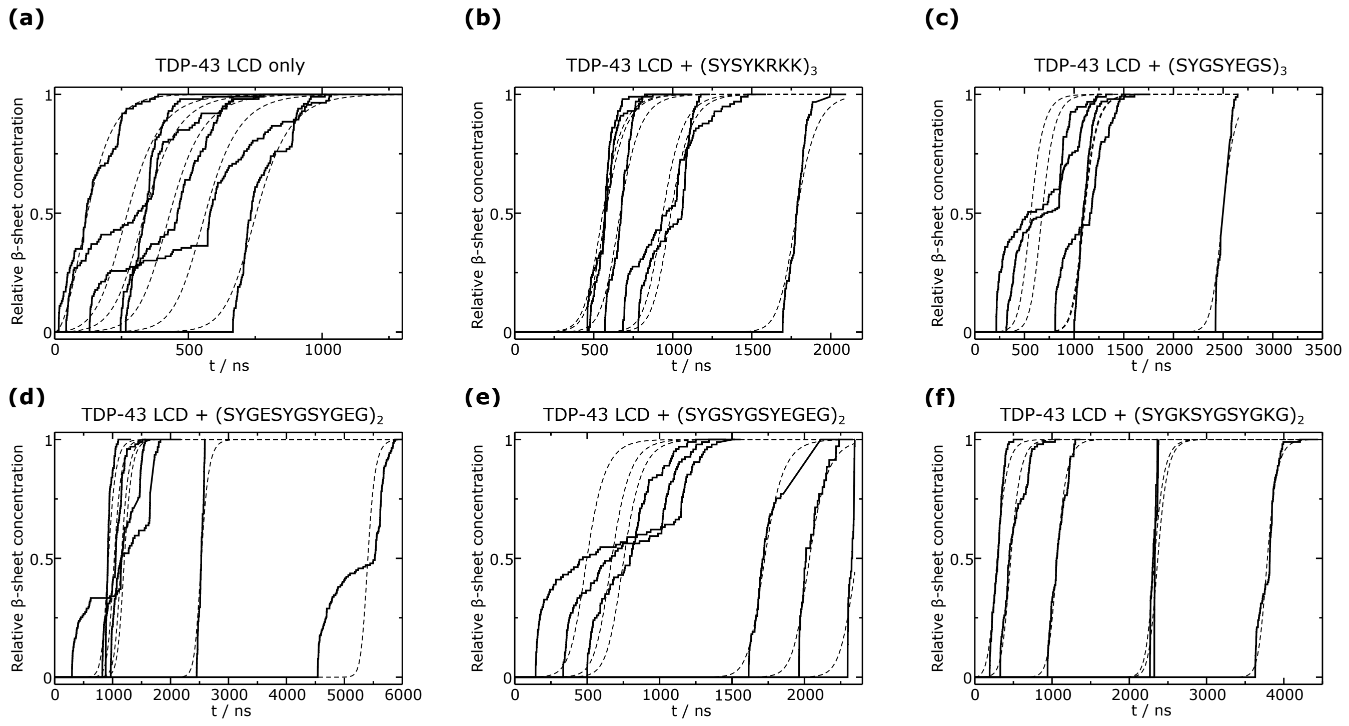


Figure S4: (a)-(f) Relative cross-β-sheet concentration for all the TDP-43 LCD systems, as indicated for each of them. For each system, the different curves represent independent trajectories. The dashed lines indicate kinetic fits performed with Amylofit software [24]. For the nature of this analysis see Section SVIII.

| | FUS LCD | | TDP-43 LCD | |
|---|---|---|---|---|
| Guest molecule | U value | p value | U value | p value |
| $(SYSYKRKS)_3$ | 7 | 0.0465 | - | - |
| $(SYSYKRKK)_3$ | 16 | 0.405 | 4.5 | 0.00415 |
| $(SYGSYEGS)_3$ | 5 | 0.0228 | 1 | 0.00415 |
| $(SYGESYGSYGEG)_2$ | 4 | 0.0154 | 0 | 0.00256 |
| $(SYGSYGSYEGEG)_2$ | 7 | 0.0465 | 2 | 0.00657 |
| $(SYGKSYGSYGKG)_2$ | - | - | 7 | 0.0467 |

Table S3: U and p values for Mann–Whitney U test comparing between the systems in presence vs. absence of the peptides.

### SIX. FREQUENCY CONTACT MAPS

As stated in the main text, the intermolecular contact frequency maps are shown here for FUS LCD and TDP-43 LCD in Figures S7 and S8, respectively. For this calculation, we consider an effective contact to take place when two residues are found at a distance below $1.2\sigma_{ij}$, being $\sigma_{ij}$ the average molecular diameter of the two residues $i$ and $j$, and 1.2 a slightly larger distance to the minimum in the potential ($\sim 1.12\sigma_{ij}$) between the $i$ and $j$ amino acids. The scale of Figure 4 (main text), S7 and S8 is in percentage of configurations in which an intermolecular contact has been found to be present.

### SX. RADIUS OF GYRATION MEASUREMENTS

As stated in the main text, we provide the radius of gyration ($R_g$) for FUS LCS and TDP-43 LCD. In Figures S9 and S10 we provide these values for the FUS LCD and TDP-43 LCD proteins respectively. The measurement are carried out in the condensate bulk, in absence and presence of the small peptides inserted in the system. We plot $R_g$ as a function of the aggregation half time ($t_{1/2}$), and we do not observe any trend or variation of the radius of gyration, suggesting that no protein conformational change is not behind the change in the aggregation kinetics.

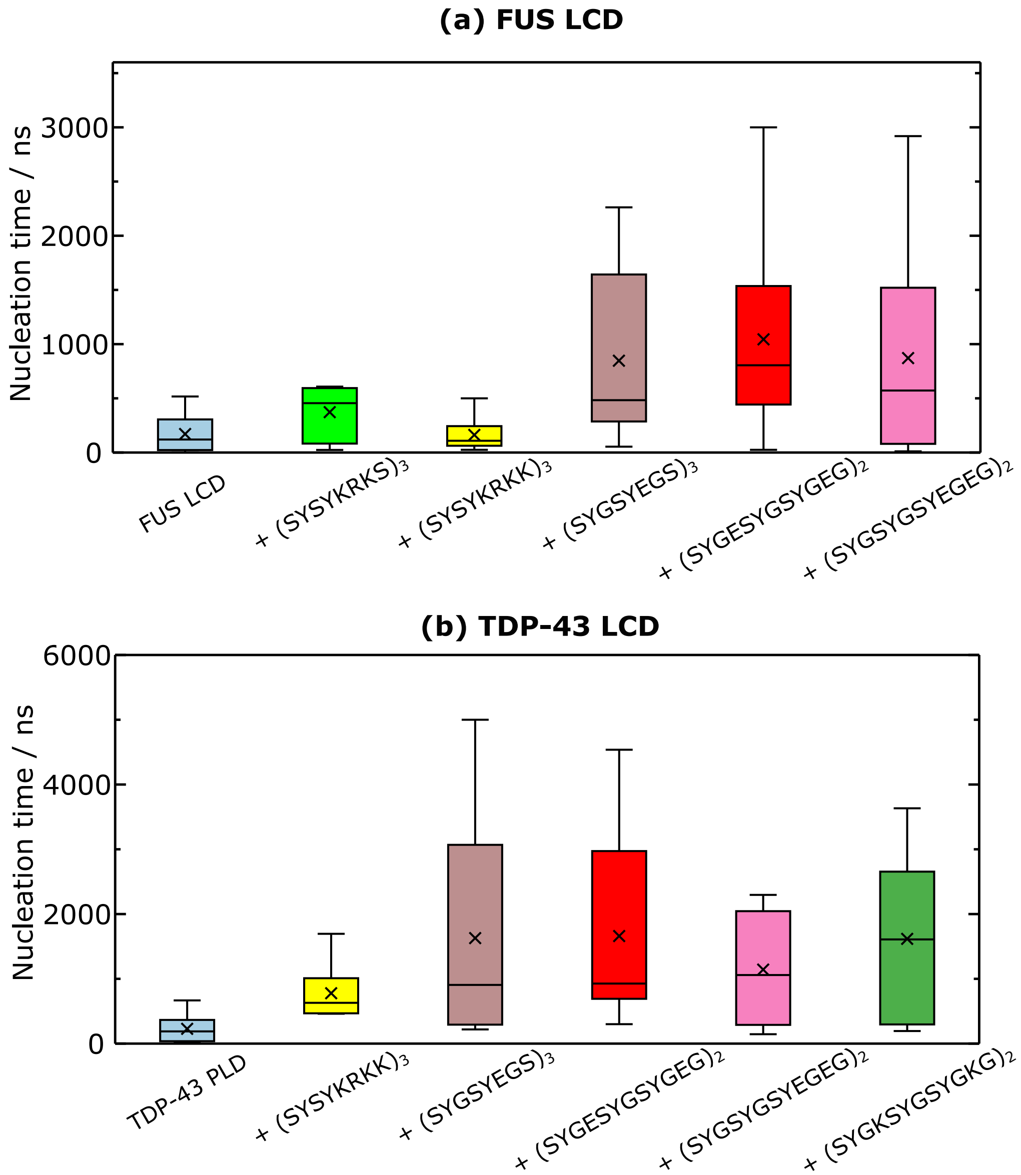


Figure S5: Nucleation time, measured as the time at which the first inter-protein β-sheet is formed, for (a) FUS LCD systems, and (b) TDP-43 PLD systems. The data is displayed as box plots, where the average value is represented with a cross symbol and the box boundaries are obtained from the 25 and 75 percentile values from our data range, exclusive.

### SXI. DIFFUSION COEFFICIENT

We measure the diffusion coefficient ($D$) of FUS LCD AND TDP-43 LCD in bulk condensate conditions. We do this both in presence and absence of the small peptides, before any inter-protein β-sheet is formed. For the proteins, we measure the mean squared displacement (MSD) of the central residue of the protein and by means of the relation MSD=6$D$t, we obtain the self-diffusion coefficient. In Figure S11 we show the results for all the systems studied throughout this work.

### SXII. (WY)$_{12}$ AGEING AND R$_G$

As mentioned in the main text, alternatively to all the system in which the density decreases, we also tested one case in which the condensate density increases upon addition of the small peptide. We choose to add (WY)$_{12}$ to FUS condensates in a concentration of 0.0782 mg (WY)$_{12}$/mg FUS LCD, and a global condensate density of 0.436 mg/cm$^{-3}$. The average nucleation time for this system is of 485 ns, which is similar to that of FUS LCD + (SYSYKRKS)$_3$, which we do not

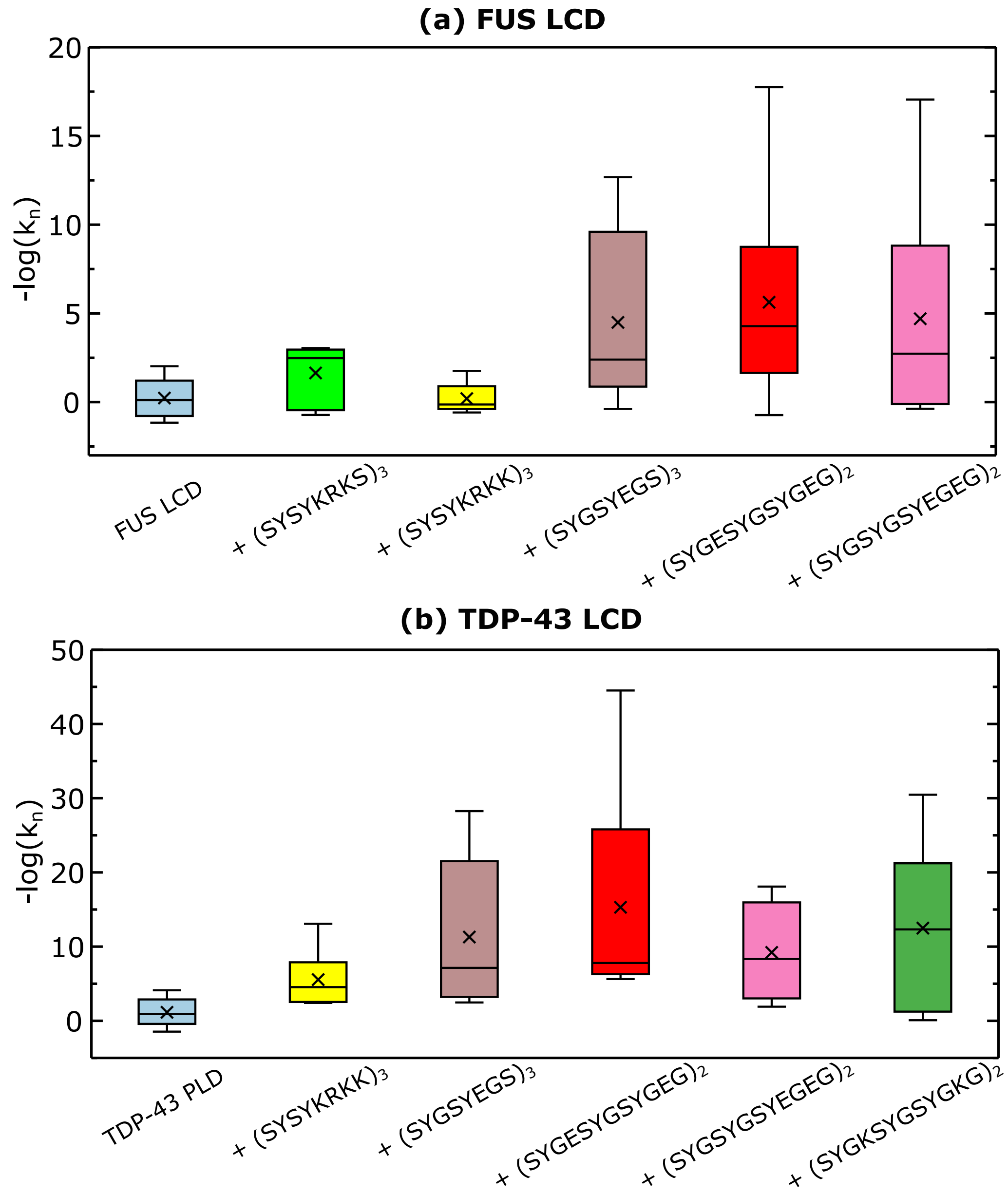


Figure S6: -log$_{10}$($k_n$) as obtained from Amylofit [24] for (a) FUS LCD, (b) TDP-43 LCD condensates, in presence of the peptides indicated in the legend. $k_2$ is in units of [relative β-sheet concentration]$^{-1}$ ns$^{-1}$.

consider to be affective ageing deceleration, yet it does not speed up the ageing kinetics despite the higher condensate density. We corroborate that this is not caused by a hindered diffusion, since the diffusion coefficient in this case is of 191 Å$^2$ns$^{-1}$, which is not significantly different from the other systems studied. We find the explanation to this intriguing behaviour in a significant change in the radius of gyration, which we measure to be 30.5(8), a decrease of about 15% in this quantity. Given that for all the other FUS LCD and TDP-43 LCD systems studied in this work the radius of gyration remains unaltered, this seems to be the crucial parameter contributing to an interesting ageing modulation. This results are reflected in Figure S12, where we compare the nucleation time and the radius of gyration for FUS LCD in pure protein condensates, as well as in presence of (SYSYKRKS)$_3$ and (WY)$_{12}$.

### SXIII. AGEING AS A FUNCTION OF CONCENTRATION

We performed non-equilibrium ageing simulations of FUS LCD with varying concentrations of peptide. We choose (SYSYKRKS)$_3$ since it is a peptide that marginally decelerates the emergence of inter-protein β-sheets. We double the

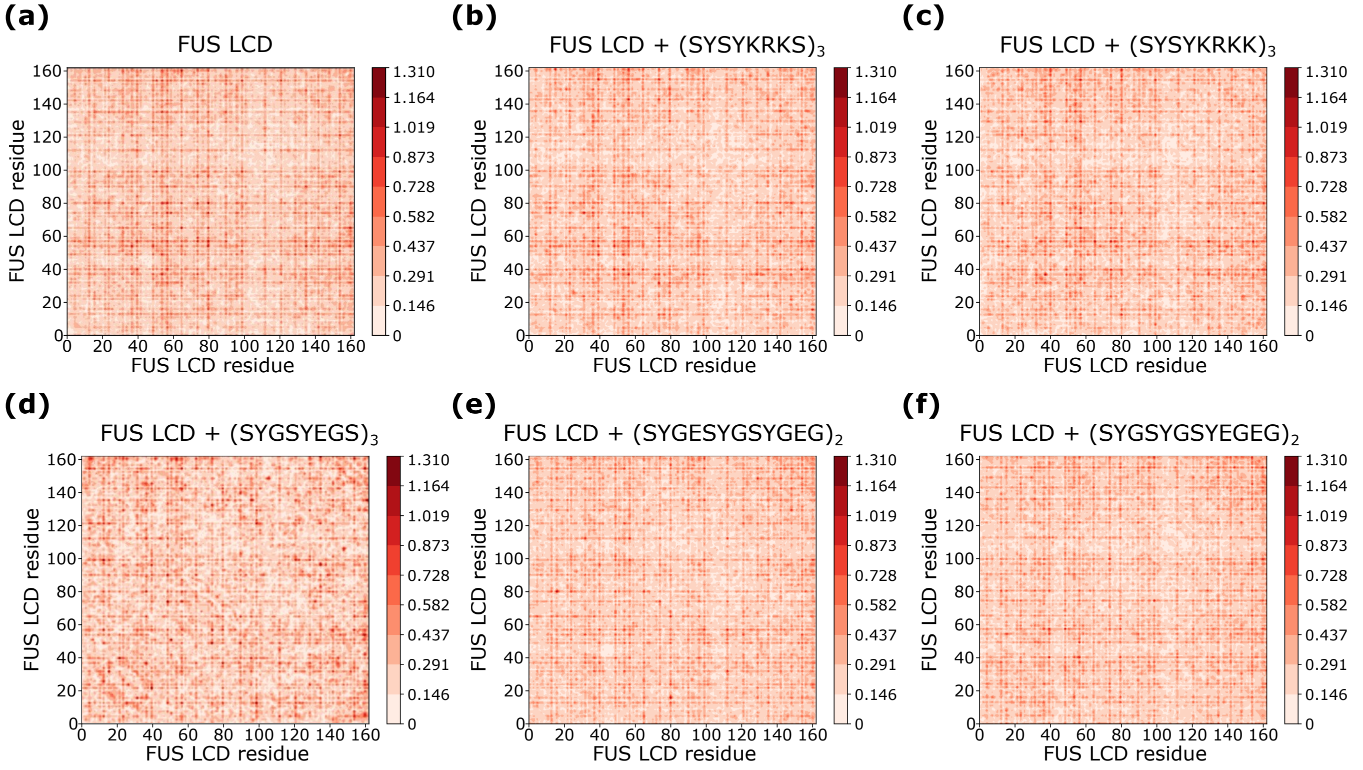


Figure S7: Intermolecular frequency contact map for the different FUS LCD systems, as indicated. The contacts in all the plots of this figure are represented as contact frequency in percentage.

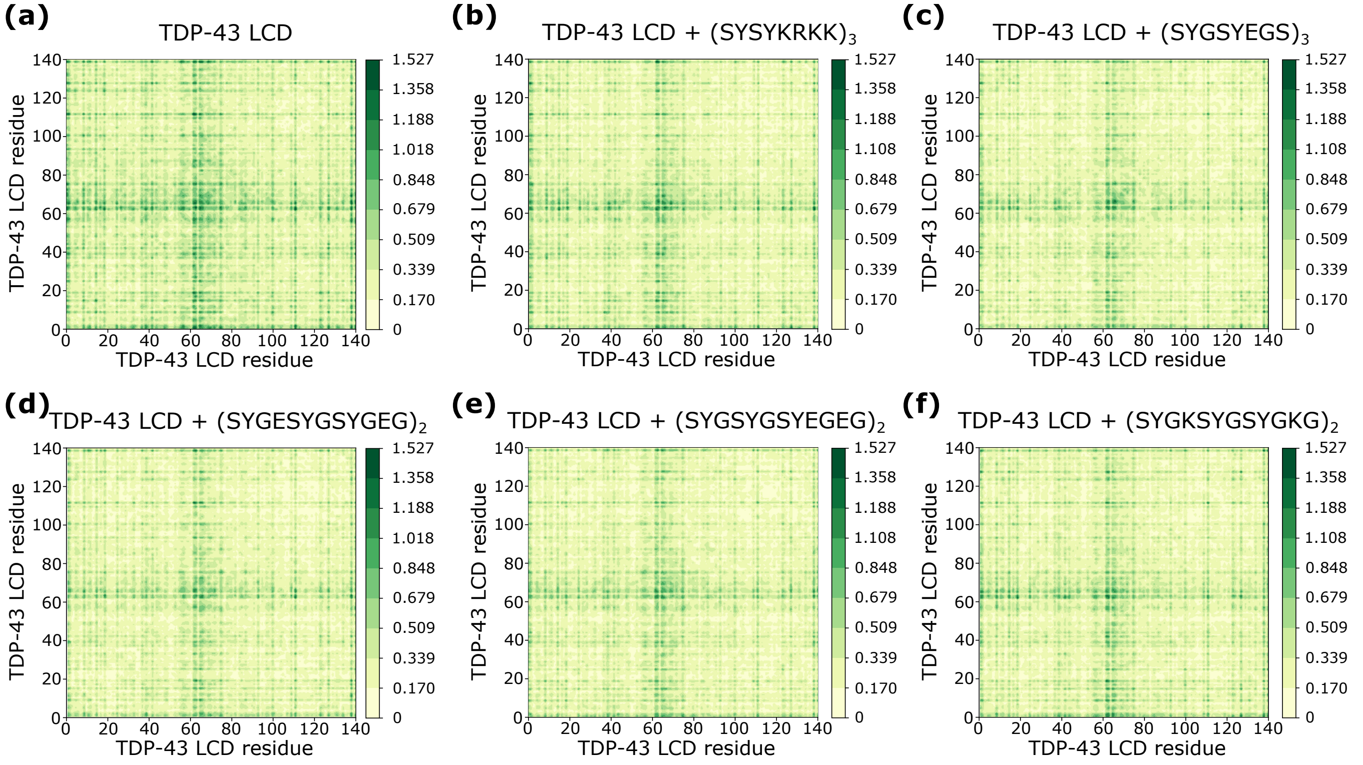


Figure S8: Intermolecular frequency contact map for the different TDP-43 LCD systems, as indicated. The contacts in all the plots of this figure are represented as contact frequency in percentage.

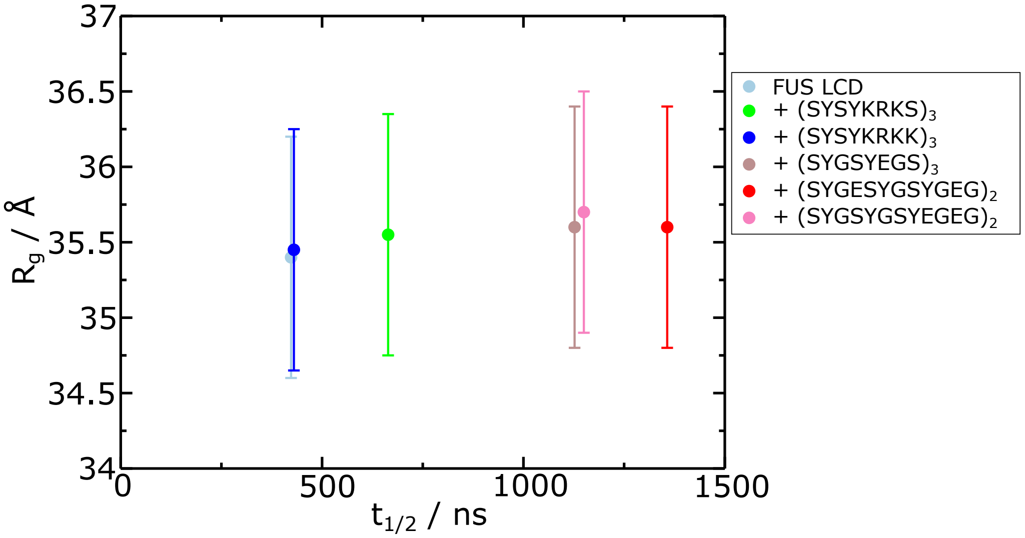


Figure S9: Radius of gyration ($R_g$) as a function of the aggregation half time ($t_{1/2}$) for FUS LCD in bulk condensate. The nature of the small peptide is specified in the figure legend.

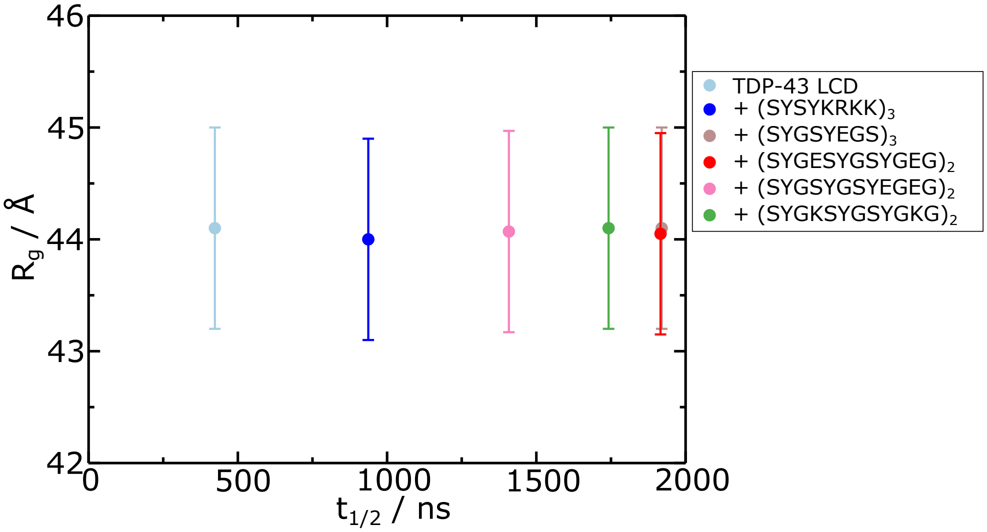


Figure S10: Radius of gyration ($R_g$) as a function of the aggregation half time ($t_{1/2}$) for TDP-43 LCD in bulk condensate. The nature of the small peptide is specified in the figure legend.

concentration of the small peptide with respect to the one that we initially selected, as explained in the main text section IIB. Under these conditions, the total concentration of peptide exceeds what we defined to be optimal, which can affect the overall mechanical properties of the condensate. In Figure S13 we show the nucleation half times for increasing concentration of $(SYSYKRKS)_3$. We observe a notable deceleration fo the ageing process with large amounts of the peptide present in the system. This behaviour is not unexpected since at such large peptide concentrations, as well as lower condensate densities (0.29 g·cm$^{-3}$), the LARKS-LARKS high density fluctuations are less probable. However, as discussed in the main text, such high peptide concentrations are out of the scope of this work since we aim to find peptides capable of disrupting the ageing process with a minimal alteration of the condensate properties. In that respect, $(SYGESYGSYGEG)_2$ and $(SYGSYGSYEGEG)_2$ perform better at decelerating ageing at lower concentrations, as shown in Figure 3(b) (main text). Figure S13 corroborates the ageing-decelerating potential of small peptides, without assessing the consequences of their insertion in the condensate.

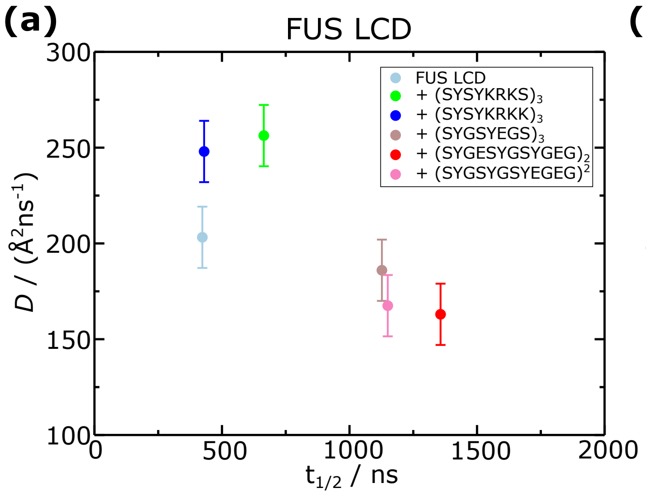


Figure S11: Diffusion coefficient for (a) FUS LCD (b) TDP-43, calculated in presence of the different small peptides studied, as indicated in the legend.

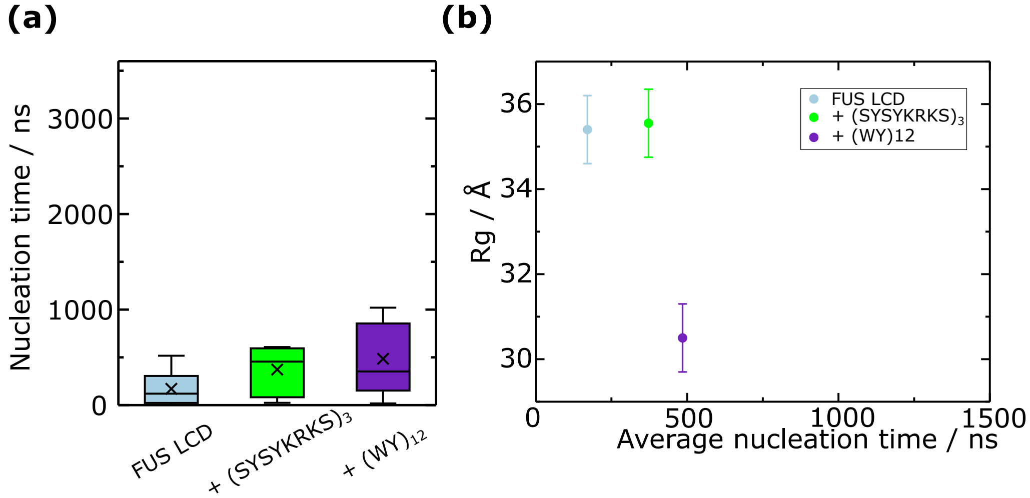


Figure S12: (a) Nucleation time represented as box plots, measured as detailed in Section SVIII for pure FUS LCD, FUS LCD + (SYSYKRKS)$_3$, and FUS LCD + (WY)$_{12}$. (b) FUS LCD radius of gyration measured from bulk simulations for the systems detailed in the legend.

### SXIV. PROTEIN SEQUENCES

FUS LCD

```
MASNDYTQQA TQSYGAYPTQ PGQGYSQQSS QPYGQQSYSG YSQSTDTSGY GQSSYSSYGQ

SQNTGYGTQS TPQGYGSTGG YGSSQSSQSS YGQQSSYPGY GQQPAPSSTS GSYGSSSQSS

SYGQPQSGSY SQQPSYGGQQ QSYGQQQSYN PPQGYGQQNQ YNS
```

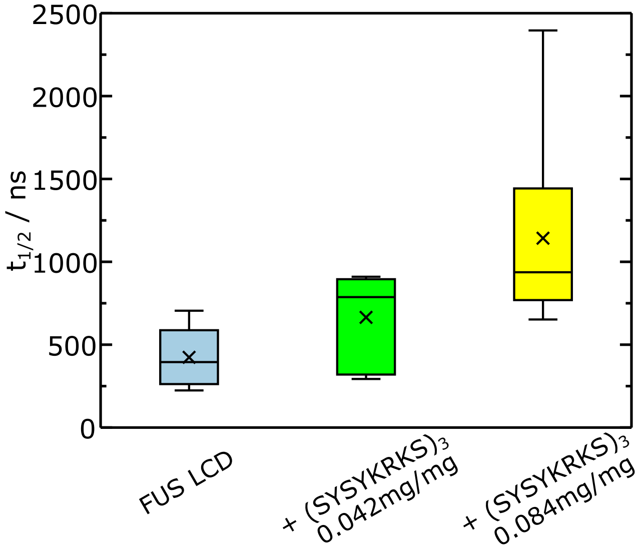


Figure S13: Nucleation half time for FUS LCD in absence and presence of the $(SYSYKRKS)_3$ peptide at varying concentrations, represented as box plots.

TDP-43 LCD

```
GRFGGNPGGF GNQGGFGNSR GGGAGLGNNQ GSNMGGGMNF GAFSINPAMM AAAQAALQSS

WGMMGMLASQ QNQSGPSGNN QNQGNMQREP NQAFGSGNNS YSGSNSGAAI GWGSASNAGS

GSGFNGGFGS SMDSKSSGWG M
```

---

[19] M. Rubinstein and R. H. Colby, *Polymer physics*. Oxford university press, 2003.

[20] A. E. Likhtman, "Single-Chain Slip-Link Model of Entangled Polymers:˜Simultaneous Description of Neutron Spin-Echo, Rheology, and Diffusion," *Macromolecules*, vol. 38, pp. 6128–6139, jul 2005.

[21] V. A. Boudara, D. J. Read, and J. Ramírez, "Reptate rheology software: Toolkit for the analysis of theories and experiments," *Journal of Rheology*, vol. 64, no. 3, pp. 709–722, 2020.

[22] J. S. Rowlinson and B. Widom, *Molecular theory of capillarity*. Courier Corporation, 2013.

[23] J. A. Zollweg and G. W. Mulholland, "On the law of the rectilinear diameter," *The Journal of Chemical Physics*, vol. 57, no. 3, pp. 1021–1025, 1972.

[24] G. Meisl, J. B. Kirkegaard, P. Arosio, T. C. Michaels, M. Vendruscolo, C. M. Dobson, S. Linse, and T. P. Knowles, "Molecular mechanisms of protein aggregation from global fitting of kinetic models," *Nature protocols*, vol. 11, no. 2, pp. 252–272, 2016.
